# Proliferation is a requirement for differentiation of oligodendrocyte progenitor cells during CNS remyelination

**DOI:** 10.1101/2020.05.21.108373

**Authors:** Sarah Foerster, Björn Neumann, Crystal McClain, Ludovica Di Canio, Civia Z Chen, Daniel S Reich, Benjamin D Simons, Robin JM Franklin

## Abstract

Oligodendrocytes that generate new myelin sheaths around demyelinated axons in the regenerative process of remyelination are generally derived from oligodendrocyte progenitor cells (OPCs). During this process, OPCs become activated, populate the area of myelin loss, and finally differentiate. Although much is known about the individual stages of remyelination, the exact sequence of events and whether a dependency of each individual stage on each other exists is unknown. Understanding the biology behind these questions is important for the development of remyelination therapies to overcome the age-related decline in remyelination efficiency observed in the chronic phase of multiple sclerosis (MS). Here we show that, following toxin-induced demyelination, all re-populating OPCs first migrate into the site of damage, undergo relatively few rounds of division and eventually differentiate. We further show that OPC proliferation is a requirement for differentiation. Together, our results reveal an unexpected link between OPC proliferation and differentiation, and opens up the possibility of novel regenerative strategies in MS focussing on stimulating OPC proliferation.

## Introduction

The central nervous system (CNS) has a remarkable capacity to regenerate new myelin sheaths in response to injury in a process called remyelination. In response to demyelination, oligodendrocyte progenitor cells (OPCs) are activated, recruited (a combination of proliferation and migration) to the site of injury, and differentiate into mature myelin-sheath-forming oligodendrocytes, eventually remyelinating denuded axons (Franklin and ffrench-Constant, 2017).

As with most regenerative processes, the efficiency of remyelination declines with age (Absinta et al., 2016; Hampton et al., 2012; Neumann et al., 2019; Pfeifenbring et al., 2015; Shields et al., 1999; Sim et al., 2002), rendering axons vulnerable for degeneration and thereby contributing to the progressive phase of multiple sclerosis (MS) (Hampton et al., 2012). Hence, remyelination-enhancing therapies are regarded as one of the pillars of future treatments of MS (Franklin and ffrench-Constant, 2017; Reich et al., 2018).

Although compromise of each of the phases of remyelination can contribute to age-related functional decline, and therefore represent potential therapeutic targets, a prevailing hypothesis is that it is the impairment of differentiation that largely accounts for remyelination failure. This hypothesis is based on experimental and neuropathological observations. First, experimentally, increasing numbers of OPCs following demyelination in an aged animal does not enhance remyelination efficiency, suggesting that the bottleneck in ageing is not the provision of OPCs but rather their ability to differentiate (Woodruff et al., 2004). Moreover, agents that enhance differentiation increase remyelination efficiency and reverse cognitive impairment in aged animals (Huang et al., 2011; Wang et al., 2020). Second, MS lesions often, although not always (Boyd et al., 2013), contain sufficient numbers of oligodendrocyte lineage cells, which seem to be arrested in their differentiation (Chang et al., 2002; Kuhlmann et al., 2008). Together, these results have led to intense efforts to identify molecules that promote OPC differentiation as potential regenerative therapies in MS.

However, despite some success in translating differentiation-based therapies into the clinic (Green et al., 2017), it is still uncertain how widely effective these will be, given that MS patients will have multiple unsynchronised lesions at different stages of their evolution. For instance, treatments that promote differentiation may disturb the recruitment phase in other lesions. Therefore, therapeutic enhancement of remyelination in MS patients will most likely need interventions that improve all phases of remyelination. The search for such interventions requires an integrated model of remyelination, including how each of the phases is related to the others and the dynamics of the OPC response throughout. Towards this goal, several important questions remain to be answered: what is the specific sequence of events that OPCs undergo during remyelination; what proportion of available OPCs contribute to remyelination; and how are the proliferation and differentiation phases related? Here, we address these questions using a widely used mouse model of toxin-induced demyelination, which allows us to follow remyelination in time and space.

## Results

To investigate the sequence of events early after lesion induction, we assessed OPC dynamics every 24 hours between 1- and 4-days post lesion (dpl) (Figure 1A). Defining the lesion boundary is critical to our analysis: at 1dpl and 2dpl it was determined by the reduction of DAPI^+^ nuclei and the absence of tissue autofluorescence, while at 3dpl and 4dpl the lesion border was identified by the higher density of DAPI^+^cells and the presence of ramified IBA1^+^ microglia in the lesion area. All sections were also stained with luxol fast blue to assist in establishing the lesion border. To label proliferating cells, while minimising the opportunity for extensive migration of labelled cells, we injected EdU 2h before sacrificing the mice. At 24 hours post lesion induction, there were no OLIG2^+^ oligodendrocyte lineage cells (OLCs) in the lesion (Figure 1B,C), suggesting that these cells underwent cell death associated with toxin injection. OLIG2^+^ OLCs were first detected in the lesion at 2dpl (Figure 1B,C), none of which were EdU^+^ (Figure 1B,D). OLIG2^+^ /^+^ells were first observed in the lesion at 3dpl (Figure 1B,D). The number of OLIG2^+^ OLCs in the lesion increased over subsequent days, while the proportion of OPCs that proliferated at a given point in time remained constant at approximately 15% (Figure 1B-D). We did not observe any OLIG2^+^ /EdU^+^ cells outside and adjacent to the lesion area at any time point (Figure 1B,D), which, together with the absence of OLIG2^+^ /EdU^+^ cells in the lesion at 2dpl, indicates that OPCs first migrate into the lesion area from surrounding intact white matter before undergoing proliferation.

**Figure 1:**
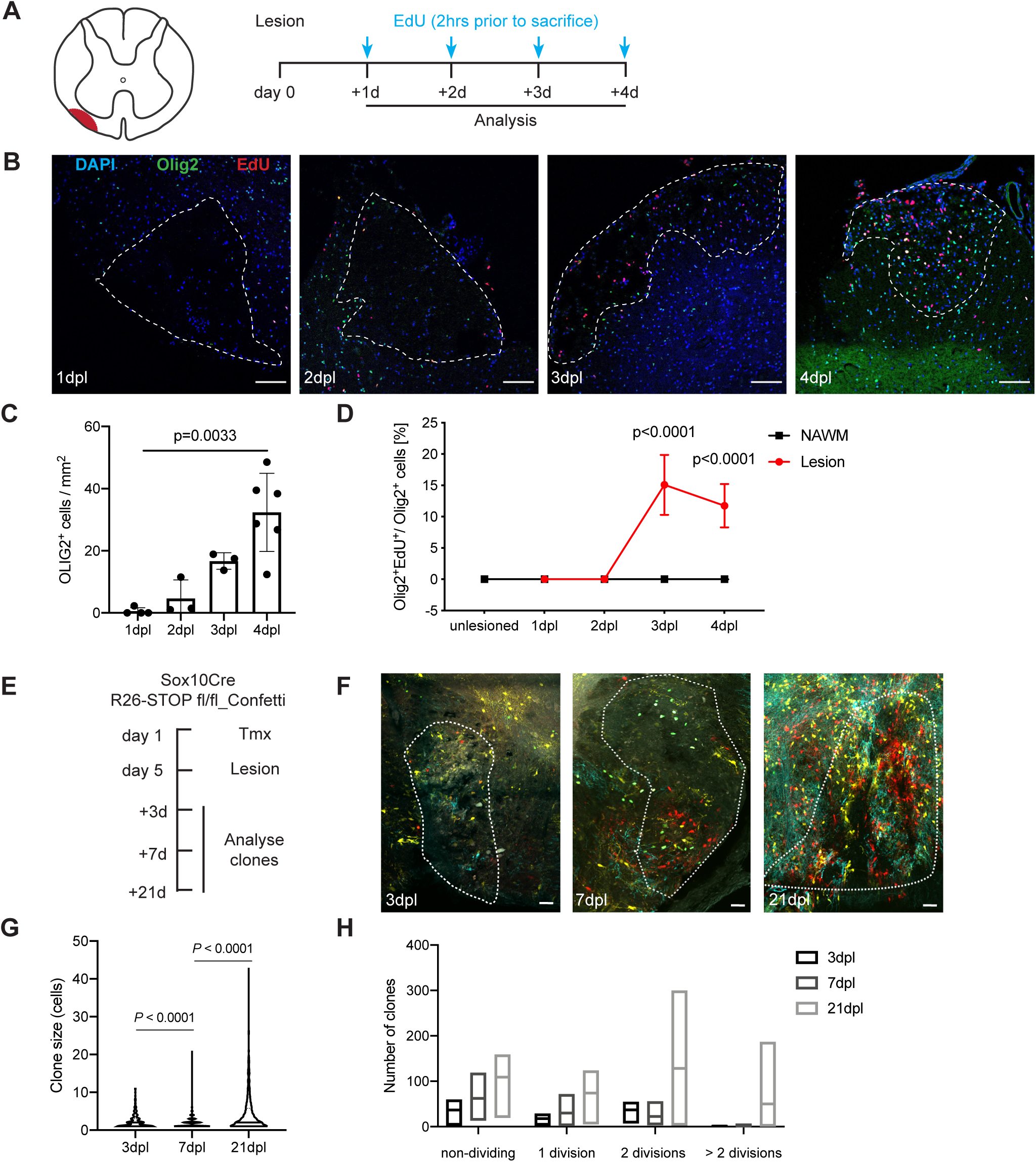
OPCs migrate into the lesion area before they proliferate. **(A)** Schematic representation of anatomical location of toxin-induced lesion in the ventral funiculus of the spinal cord (left). Experimental outline (right). d = day **(B)** Representative images of immunohistochemistry staining of lesions. dpl = days post lesion. White line denotes the lesion area. Scale bar: 100*µ*m. **(C)** The number of OLIG2^+^ cells per lesion area (Kruskal-Wallis test). **(D)** Percentage of proliferating EdU^+^ OLIG2^+^ cells in the normal appearing white matter (NAWM) and lesion area (2-way ANOVA). **(E)** Experimental outline. Tmx = Tamoxifen. **(F)** Representative images of oligodendrocyte lineage cell clones in Sox10-Cre-confetti mouse. White line denotes the lesion area. Scale bar: 25*µ*m. **(G)** Violin plots depicting the size of individual clones at different time points after lesion induction. Kruskal-Wallis test followed by Dunn’s multiple comparison test. **(H)** Number of cell divisions undergone by individual clones, which was calculated from the total number of cells in one clone (2-way ANOVA) (n(7dpl)=228, n(14dpl)=583 and n(21dpl)=1452 clones).

To address what proportion of peri-lesion OPCs contribute to the repopulation of the lesion, we used a tamoxifen inducible *Sox10iCre*-driven confetti reporter mouse to follow individual OPC clones responding to lesion induction (Figure 1E) (Laranjeira et al., 2011; Snippert et al., 2010). Since lysolecithin led to the ablation of Sox10^+^ oligodendrocyte lineage cells within the lesion, only invading OPCs can have contributed to the formation of clones within the lesions (Figure 1C). To distinguish individual OLC clones, we used hierarchal clustering of fluorescent cells within the lesion. To obtain a 3D representation of the lesion, three serial sections, each 100*µ*m thick, were stacked. We found that the number of clones in the lesion increased with time (Figure 1F,G), indicating that there is sustained migration of SOX10^+^ OLCs into the lesion. The majority of SOX10^+^ clones contained less than 5 cells, indicating 3 or fewer divisions (Figure 1H). The number of divisions of OPCs can be inferred from the clone size since very few OLCs undergo cell death at early stages of the lesion (Ruckh et al., 2012). Only at 21dpl were clones comprising more than 8 cells (i.e > 3 divisions) observed (Figure 1H), indicating expansion of at least some clones. Together, these data suggest a model of lesion repopulation involving many cells responding to demyelination by first migrating into the lesion and there undergoing relatively few cycles of cell division.

The final phase of remyelination is when recruited OPCs differentiate and mature into non-dividing, myelin sheath-forming oligodendrocytes. However, whether OPC proliferation is necessary for oligodendrocyte differentiation is unclear. To address this, we labelled all proliferating OPCs by long-term administration of EdU and then assessed the proportion of cells expressing the differentiation-associated maker CC1 that were EdU^+^ (Figure 2A). At 14 dpl, we found that the vast majority of CC1^+^ oligodendrocytes (96%) were also labelled with EdU (Figure 2B,C). Since CC1^+^ cells do not divide in this model (Crawford et al., 2016), this indicates that CC1^+^ cells are derived from cells that have undergone proliferation (Figure 2C).

**Figure 2:**
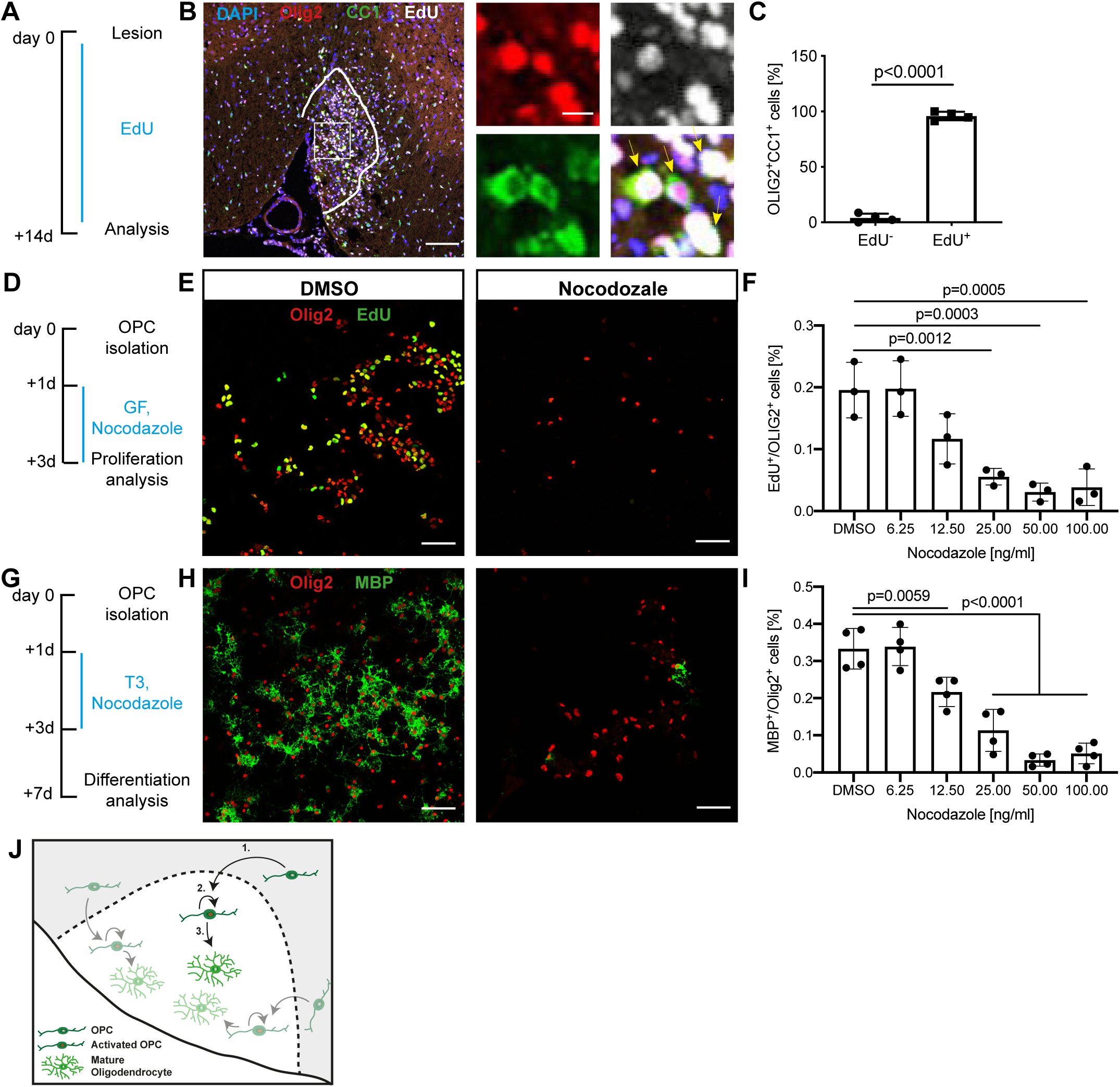
OPCs proliferate before they differentiate. **(A)** Experimental outline of Fig 2B,C. d = day **(B)** Representative images of immunohistochemistry staining of lesions at 21 days post lesion. White line denotes the lesion area. Arrowheads mark EdU^+^ mature CC1^+^ oligodendrocytes. Scale bar: 100*µ*m and 10*µ*m. **(C)** Percentage of mature CC1^+^ oligodendrocytes that are EdU^+^ due to prior OPC proliferation (unpaired t-test). **(D)** Experimental outline of Fig 2E,F. GF = growth factors. **(E)** Representative images of immunocytochemistry staining of OPCs treated with DMSO or Nocodazole after 3 days in culture. Concentration nocodazole: 50ng/ml. Scale bar: 100*µ*m. **(F)** Proportion of EdU^+^ OLIG2^+^OPCs following DMSO or nocodazole treatment at various concentrations (1-way Anova). **(G)** Experimental outline for Fig 2H,I. **(H)** Representative images of OPC-derived oligodendrocytes treated with DMSO or nocodazole at 7 days in culture. Concentration nocodazole: 50ng/ml. Scale bar: 100*µ*m. **(I)** Proportion of MBP^+^ OLIG2^+^ oligodendrocytes following DMSO or nocodazole treatment at various concentrations (1-way ANOVA). **(J)** Summary of the sequence of events that OPCs undergo in response to toxin-induced lesions.

To test if proliferation is required for OPC differentiation, we blocked cell cycle progression in OPCs in a reversible and non-toxic way during differentiation using nocodazole, a M-phase inhibitor (Yiangou et al., 2019). Treatment with nocodazole for 48h blocked the division of neonatal OPC cultured in a proliferation medium in a dose-dependent manner (Figure 2E,F). Since no significant reduction in OPC numbers per well was observed in response to nocodazole treatment, even at a concentration of 100ng/ml (Figure S1), we concluded that nocodazole treatment was effective and non-toxic below this dose. We then cultured OPCs in differentiation medium, which included T3. To block mitosis, we used several concentrations of nocodazole for the first 48h of differentiation. This resulted in a significant reduction in the differentiation of OPCs into mature MBP^+^ oligodendrocytes after 5 days (Figure 2G-I), consistent with the conclusion that proliferation is directly linked to differentiation. Alternatively, this result could arise from the majority of OPCs committing to self-renewal, having passed the G1/S checkpoint, at the beginning of the experiment. To distinguish between these possibilities, we assessed the cell cycle phase profile of OPCs after isolation using flowcytometry analysis after staining with propidium iodine. We found that 60% of primary isolated OPCs are in G0/G1 (Figure S2). Since most stem and progenitor cells undergoing terminal differentiation make a cell fate decision to undergo either proliferation or differentiation in G1-phase of the cell cycle (Soufi and Dalton, 2016), it is likely that the OPCs were competent to directly undergo differentiation when we started the treatment with nocodazole. We therefore concluded that OPCs need to enter the cycle before undergoing differentiation.

## Discussion

Here we show that all OPCs, albeit asynchronously, undergo the same sequence of events following experimentally-induced demyelination (Figure 2J). Initially, OPCs migrate into the site of injury and subsequently proliferate (Figure 1). Proliferation of OPCs was only observed inside the lesion and not in the non-lesion area surrounding the lesion (Figure 1). This suggests an injury-specific mode of proliferation that is different from the contact-inhibition model of proliferation observed in homeostasis (Hughes et al., 2013; Rosenberg et al., 2008). The contact-inhibition model proposes that OPC proliferation is inhibited by contact via their processes. If OPCs die or differentiate, the lack of contact inhibition induces proliferation of surrounding OPCs, re-establishing the *status quo*. Applied to the situation of large focal demyelination, this model predicts that OPC loss within the lesion would first trigger proliferation of OPCs at the lesion’s edge and subsequent migration of one or even both daughter cells into the lesion, the latter event triggering further proliferation within non-lesioned tissue. However, this is not what was observed in our study. Instead, our data suggest that the first OPC response to demyelination is migration into the lesion, likely from a relatively narrow zone (Franklin et al., 1997), where they subsequently undergo proliferation. This model is consistent with observation that migration of OPCs into the lesion is mainly restricted to the first days following demyelination (Niu et al., 2019).

Following proliferation, OPCs differentiate into mature oligodendrocytes (Tripathi et al., 2010; Zawadzka et al., 2010). In this study, most oligodendrocytes arose from OPCs that have previously undergone proliferation (Figure 2), similar to OPCs in the developing zebrafish CNS (Marisca et al., 2020). However, this is at variance with the study by Hughes and colleagues (Hughes et al., 2013), who found that in only the minority of events (7 out of 107 cells) did proliferation precede differentiation in the adult cortex. The possible reasons for this discrepancy between the studies are numerous and include the experimental paradigm (homeostasis versus injury) and CNS region (grey matter in the brain versus white matter in the spinal cord). Our data indicate that proliferation may be required for differentiation following an *in vivo* injury. Indeed, we showed in tissue culture experiments that blocking proliferation in neonatal OPCs significantly reduced their differentiation capacity. The changes in OPC density, resulting from the block of proliferation, could hypothetically have also contributed to differentiation failure. However, we have recently shown that halving the cellular density per well, does not alter differentiation capacity of OPCs (Neumann et al., 2019). Together, our data suggest that proliferation is directly linked and required for differentiation *in vitro* and during remyelination *in vivo*.

This finding has important implications for the understanding of why OPC differentiation, and thus remyelination, is impaired in the setting of age and disease. A prevalent hypothesis to explain the decline in remyelination efficiency with age is a failure of OPCs to differentiate into mature oligodendrocytes in a timely fashion (Boda et al., 2015; Shen et al., 2008; Woodruff et al., 2004). Recently, we have shown that the exposure of aged OPCs to differentiation-enhancing compounds cannot enhance their differentiation, suggesting that aged OPCs cannot respond to differentiation signals (Neumann et al., 2019). Our finding that OPCs must undergo proliferation to differentiate might offer an intriguing explanation for this observation since aged OPCs have a markedly reduced capacity to proliferate both *in vitro* and *in vivo* (Ruckh et al., 2012; Segel et al., 2019). According to this hypothesis, interventions that can induce proliferation might also restore the differentiation capacity of aged OPCs. We have recently demonstrated that soft hydrogels, mimicking the stiffness of a neonatal brain, can restore not only the proliferation, but also the differentiation capacity of aged OPCs. Further, we have found that the overexpression of *MYC*, an oncogene, causes aged OPCs to re-enter the cell cycle and subsequently undergo differentiation at levels comparable to young, proliferation competent OPCs (Neumann et al., 2020, under review). Similar effects are observed in other stem cell systems that have comparable dynamics to OPCs. For example, post-mitotic juvenile cardiomyocytes can be induced to proliferate by the expression of Myc (Bywater et al., 2020), while aged muscle stem cells can be rejuvenated by activation of Notch signalling, an intervention that restores their proliferation capacity and concomitantly allows for efficient differentiation of these cells (Conboy et al., 2003; Liu et al., 2018). Equally, neural stem cells become increasingly quiescent with age and stop generating new functional neurons, a process that is linked to increased inflammation and reduced mitogens, both compromising proliferation (Kalamakis et al., 2019). Therefore, there is strong evidence in our and other studies that reinstating proliferation capacity is necessary for the restoration of regenerative function of several adult stem cell systems.

The question that follows is whether the strategies for remyelination enhancement should shift to focus on interventions that mobilise potentially quiescent OPCs into proliferation, a state of activation that will then allow differentiation. Although our results indicate that differentiation is intrinsically tied to proliferation, there is no direct evidence that exposure of proliferation stimulating agents may resolve the age-related, and disease relevant, decline in remyelination efficiency. Aged OPCs cultured in increasing amounts of PDGF-AA and FGF2 fail to proliferate even after extended periods of time *in vitro* (Segel et al., 2019). Thus, we suggest that it is the diminished intrinsic stem cell potential to enter self-renewal which underlies the impairment of aged OPCs to remyelinate. Therefore, interventions that re-activate the stem cell potential and thus the proliferation capacity of aged OPCs might be suitable interventions to overcome the age-related decline in remyelination efficiency in patients with MS.

## Acknowledgments

This work was supported by funding from the UK Multiple Sclerosis Society (MS 50), the Adelson Medical Research Foundation, the Intramural Research Program of NINDS and a core support grant from the Wellcome and MRC to the Wellcome-Medical Research Council Cambridge Stem Cell Institute (203151/Z/16/Z). We thank the NIHR Cambridge BRC Cell Phenotyping Hub for all flow cytometry work. B.D.S acknowledges funding from the Royal Society (E.P. Abraham Research Professorship, RP\R1\180165) and Wellcome Trust (098357/Z/12/Z).

## Author Contributions

Conceptualization, S.F., B.N and R.J.M.F.; Methodology, S.F., B.N and C.R.M.; Investigation, S.F., B.N., C.R.M., L.D.C.; Analysis and Interpretation, S.F., B.N., C.R.M., L.D.C., C.Z.C., D.S.R., B.D.S. and R.J.M.F.; Writing S.F., B.N., C.R.M. and R.J.M.F.; Funding Acquisition B.D.S. and R.J.M.F.; Supervision B.D.S. and R.J.M.F.

## STAR methods Text

### Lead contact and materials availability

Further information and requests for resources and reagents should be directed to and will be fulfilled by the Lead Contact, Robin JM Franklin. This study did not generate new unique reagents.

### Experimental models and subject details

#### Animal husbandry

All animal procedures were performed in compliance with United Kingdom Home Office regulations. The animals were housed under standard laboratory conditions on a 12 h light/dark cycle with constant access to food and water. All animals were housed in pairs or groups of up to 6 animals. All animals were randomly allocated to experimental groups. All experimental groups were matched for age and sex. Detailed information for the source of all rat and mouse strains is available in the KRT.

#### Isolation of neonatal oligodendrocyte progenitor cells

Neonatal (P5-7) rats were decapitated after lethal injection with phenobarbital. The brains were removed quickly and placed into ice-cold isolation medium (Neumann et al., 2019) (alternatively Hibernate A Brainbits). The telencephalon and cerebellum were dissected in isolation medium; meninges, and the olfactory bulb were mechanically removed and the brain tissue was mechanically minced into 1mm^3^ pieces. The tissue pieces were spun down at 100g for 1min at room temperature (RT) and the tissue was washed in HBSS- (no Mg2+ and Ca2+, Gibco). Each half of the brain was mixed with 5ml of dissociation solution (34U/ml papain (Worthington), 20μg/ml DNAse Type IV (Gibco) in isolation medium). The brain tissue was dissociated on a shaker (50rpm) for 40 min at 35 ºC. The digestion was stopped by addition of ice cold HBSS-. The tissue was centrifuged (200g, 3 min, RT), the supernatant completely aspirated and the tissue resuspended in isolation medium supplemented with 2% B27 and 2mM sodium-pyruvate (trituration solution). The tissue was allowed to sit in this solution for 5min. To obtain a single cell suspension the tissue suspension was triturated 10 times using first a 5ml serological pipette and subsequently three fire polished glass pipettes (opening diameter >0.5mm). After each trituration step the tissue suspension was allowed to sediment (approximately 1-2 min) and the supernatant (approximately 2ml), containing the cells, was transferred into a fresh tube. After each round of trituration 2ml of fresh trituration solution were added. To remove accidentally transferred undigested tissue bits, the collected supernatant was filtered through 70μm cell strainers into tubes that contained 90% isotonic Percoll (GE Healthcare, 17-0891-01, in 10xPBS pH7.2 (Lifetech). The final volume was topped up with phenol-red free DMEM/F12 with HEPES (Gibco) and mixed to yield a homogenous suspension with a final Percoll concentration of 22.5%. The single cell suspension was separated from remaining debris particles by gradient density centrifugation (800g, 20min, RT, without break). The myelin debris and all layers without cells were discarded and the brain cell containing phase (last 2ml) and cell pellet were resuspended in HBSS^+^ and combined in a fresh 15ml tubes and centrifuged (300g, 5min, RT). The cell pellet was resuspended in red blood cell lysis buffer (Sigma, R7757) and incubated for 90s at RT to remove red blood cells. 10ml of HBSS+ were added to this cell suspension and spun down (300g, 5min, RT). The cell pellets were resuspended in 0.5ml modified Miltenyi washing buffer (MWB, 2mM EDTA, 2mM Na-Pyruvate, 0.5% BSA in PBS, pH 7.3) supplemented with 10ng/ml human recombinant insulin (Gibco). To this cell suspension 2.5*µ*g mouse-anti-rat-A2B5-IgM antibody were added for every 10 million cells. After 25 min incubation, gently shaking at 4 ºC, 7ml of MWB were added. The solution was centrifuged (300g, 5min, RT) and the pellet resuspended in 80μl MWB supplemented with 20μl rat-anti-mouse-IgM antibody (Milteny, 130-047-302) per 10 million cells. The cells were incubated for 15 min, slowly shaking at 4 ºC. The secondary antibody was again washed out with 7ml MWB and the sample was centrifuged (300g, 5min, RT). The cell pellet was resuspended in 0.5ml and MACS was performed according to the recommendations of the supplier. Briefly, a MS column (Milteny, 130-042-201) was inserted into MiniMACS Separator (Miltenyi; 130-042-102) and pre-wet with 0.5ml MWB. Resuspended cells were put onto one MS column. Subsequently the column was washed three times using 500μl MWB for each wash. Finally, A2B5 positive cells were flushed out the column with 1ml pre-warmed, CO2 and O2 pre-equilibrated OPC medium.

#### Flow cytometry analysis of primary OPCs

Neonatal OPCs were plated in T75 flasks and left to recover overnight. The cells were then dissociated by removing the media and incubating the cells with 1ml of 1X TrypLE (Thermofisher) for 6min at 37°C. The cells were pipetted up and down to fully detach them from the flask, transferred to a falcon tube and were spun at 300g for 5min. The cells were then transferred to low binding microcentrifuge tubes. The cells were stained with a live/dead stain by incubating them with Zombie Violet (1:100, Biolegend) in 1X PBS for 15min at room temperature in the dark. They were washed once with 1X PBS and then fixed using 70% EtOH in 1X PBS for 2hr at 4°C. The OPCs were then washed with 1X PBS and were incubated with a FxCycle™ PI/RNase Staining Solution (Thermofisher) for 30min at 37°C. The cells were then transferred to flow tubes and cell cycle stages were assessed using a BD Fortessa instrument equipped with 405nm and 561nm lasers. Results were analysed using the ‘cell cycle analysis’ feature of FlowJo.

#### Culture of neonatal oligodendrocyte progenitor cells

Isolated OPCs were seeded into 96 well-plates (InVitro-Sciences) coated with PDL (Sigma). After isolation, OPCs were left to recover in OPC medium (60*µ*g/ml N-Acetyl cysteine (Sigma), 10µg/ml human recombinant insulin (Gibco), 1mM sodium pyruvate (Gibco), 50µg/ml apo-transferrin (Sigma), 16.1µg/ml putrescine (Sigma), 40ng/ml sodium selenite (Sigma), 60ng/ml progesterone (Sigma), 330µg/ml bovine serum albumin (Sigma)) supplemented with b-FGF and PDGF (30ng/ml each, Peprotech). OPCs were incubated at 37 ºC, 5% CO2 and 5% O2. The medium was completely exchanged to OPC medium with 20ng/ml bFGF and PDGF after overnight culture to remove any dead cells. After 3d the cell culture medium was switched to promote further proliferation (OPC medium+20ng/ml bFGF and PDGF) or differentiation (OPCM + 40ng/ml T3). During differentiation or proliferation experiments 66% of the medium was replaced every 48h and growth factors or other small molecules were added fresh to the culture. The culture medium used was 100μl for cultures in 96 well plate wells. For proliferation and differentiation assays the medium was in some instances supplemented with 40ng/ml thyroid-hormone (T3, Sigma) and varying concentrations of nocodazole (Sigma, M1404).

### Method details

#### Induction of white matter lesions and assessment of remyelination

A focal demyelinating spinal cord lesion was induced in 8-12-week-old C57 Black 6 mice as previously described (Woodruff et al., 2004). The mice were anesthetized with buprenorphine (0.03mg/kg, s.c.) and 2.5% isoflurane. The spinal cord was exposed between two vertebrae of the thoracic column. Demyelination was induced by injection of 0.8µl of 1% lysolecithin (L-lysophosphatidylcholine, Sigma) into the ventral funiculus at a rate of approximately 1 µl/min. After lysolecithin delivery, the injection needle remained in position for an additional minute.

#### Immunofluorescence for tissue sections

Mice received a lethal dose of pentobarbitol and were transcardially perfused with 4% paraformaldehyde (PFA) in PBS. The brains were removed and post-fixed for 2h at RT with 4% PFA. After a rinse in PBS the tissue was incubated in 20% sucrose solution (in PBS) overnight. The tissue was then embedded in OCT-medium (TissueTek) and stored at -80ºC. 12 μm sections were obtained using a cryostat. Tissue sections were air dried and stored at -80 ºC. Cryostat cut sections were dried for 45 min at RT. Tissue sections were incubated with blocking solution (10% [v/v] normal goat serum and 0.1% [v/v] Triton X-100 in PBS] for 1 hr at 20°C–25°C. In case antibodies raised in mouse were used, tissue sections were incubated with the MOM kit according to manufacturer’s protocol (Vector laboratories). In case the antigen required antigen retrieval for detection, slides were placed in antigen retrieval buffer solution (Dako, S2369, diluted 1:10 in distilled H2O) which was preheated to 95°C and slides were incubated for 10 min at 75°C. Subsequently, tissue sections were incubated with primary antibodies overnight at 4°C, and then secondary antibodies for 1 hr at 20°C–25°C.

Cell nuclei were visualized by staining with Hoechst 33258 (1:10000, Biotium). All images were acquired using an SP5 confocal microscope (Leica). Cell counts were obtained from 3-4 biological replicates, 3-4 tissue sections per biological replicate. The lesion border was determined by the reduction of DAPI^+^ nuclei and the absence of tissue autofluorescence in the lesion area at 1dpl and 2dpl. In contrast, at 3dpl and 4dpl the lesion border was identified by the higher density of DAPI^+^ cells and the presence of ramified IBA1^+^ microglia in the lesion area. ImageJ (Version 2.0.0-rc-68/1.52h), Cell Profiler (Version 2.2.0) and Cell profiler analyst (Version 2.2.1) were used for image analysis, cell counting and measurement of the size of the selected area.

#### Immunofluorescence for cells

Cultured cells were rinsed with PBS before fixation with 4% PFA (10 min, RT). Subsequently, the cells were washed three times with PBS (5 min, RT, shaking). If permeabilisation was required, the cells were incubated with PBST (0.1% Triton-X-100 in PBS) for 10 min at RT. The samples were then blocked in PBS supplemented with 10% normal donkey serum (NDS). Primary antibodies were diluted in PBS with 5% NDS and incubated overnight at 4 ºC in a humidified chamber. Excess antibodies were washed off with three washes in PBS (10 min, RT, shaking). The primary antibodies were then labelled with secondary antibodies diluted in PBS with 5% normal donkey serum. Again, excess antibody was washed off with three washes PBS (10 min, RT, shaking). If visualisation of nuclei was required the first wash contained 2μg/ml Hoechst 33342 (Sigma). Images were taken with a Leica-SP5 (Leica) or Leica-SP8 (Leica) microscope. For 96 well plate assays cells were kept in PBS after staining. Further image processing and analysis was performed using the ImageJ software package (Schindelin et al., 2012).

#### EdU incorporation assays, immunofluorescence, and imaging

For the EdU incorporation assay, 10µM EdU was added into the cell culture medium for 2hr, followed by the protocol provided by the Click-iT Plus EdU Alexa Fluor 647 Imaging Kit (Thermo Fisher; C10640). Otherwise, *in vitro* tissue culture cells were fixed in 4% paraformaldehyde (Thermo Fisher; 10131580) for 10min at room temperature. The cells were then washed once in PBS (Thermo Fisher; BP3994) and then blocked in PBS with 0.1% Triton X-100 (Sigma; T8787) and 5% Donkey Serum (Sigma; D9663) for 30min at room temperature. EdU detection was then performed by the protocol provided with the kit and immunostaining was performed subsequently as described above. For EdU incorporation assays in vivo, mice were injected i.p. with EdU (75µg/g of bodyweight in DMSO and sterile saline) 2h before perfusion fixation with PFA as described above. Alternatively, EdU was supplemented in the drinking water (0.2mg/ml) starting from one day after the lesion, and was replaced on Monday, Wednesday and Friday every week. EdU detection was carried out according to the manufacturer’s protocol and immune staining was performed subsequently as described above. Fixed, fluorescent cells or slides were imaged using the Zeiss Axio Observer or the Leica TCS SP5 confocal microscope.

#### In vivo clonality experiments

Sox10::iCre, ROSA26::CAG-confetti mice received one tamoxifen injection i.p. (55mg/kg bodyweight). The mice were lesioned as described and sacrificed and perfused with 4%PFA at the indicated time points. Tissues were embedded in low melting point agarose and sliced into 100µm section using a vibratome. DNA in the samples was stained using TOPRO3 DNA dye and 100µm z-projections were acquired using a Leica-SP-5 with HyD detectors.

### Quantification and statistical analysis

#### Confetti quantification and clonal analysis

Fluorescent cells were mapped, and serial sections were stacked in 3D using Neurolucida software, then exported for further analysis in MATLAB (v. R2016a). The mapping coordinates were imported into structures according to fluorescent expression with our Nldata2struct MATLAB script (S1). In order to adjust for shrinkage during tissue processing and imaging, the MATLAB script ShrinkageCorrection (S2) adjusted for optical Z section to the physical cut section Z depth, with the expansion factor (EF) being manually adjusted for each animal’s section sample size. This script outputs expanded structures for corrected measurements en mass or per colour combination. The sample size within each mouse is important in determining total cluster volume, size, range, and recombination rate. It is more reliable to measure mosaically labelled cluster expansion sizes within an animal, compared to between-mouse variation using irregular cluster cross sections. The number of mosaic-labelled clusters was normalized to the total number of clusters detected in each mouse before analysis to adjust for the cluster size frequency within a mouse in order to better compare between mice. The MATLAB script Renormalisation (S3) calculated the nearest neighbours for each recombination type and divided this into bins over the area and renormalized the counts in each bin by the bin size, as previously performed to calculate mosaic recombination events. Hierarchical clustering was used as an unbiased algorithm following statistical probabilities for grouping cells of the same Confetti gene expression using nearest neighbour calculations (MATLAB script HierarchicalClusteringBoundaryPlot).

#### Statistical analysis

All statistical analysis was performed in GraphPad Prism (GraphPad Software, Inc.). Prior to the analysis of parametric data, we performed Shapiro-Wilk tests to exclude non-normality of the data. For data derived from the quantification of immunohistochemical staining, comparisons between two groups were performed with an unpaired t-test assuming two-tailed distribution and equal variances. In cases where more than two groups were compared to one-another, a one-way analysis of variance (ANOVA) was performed assuming equal variances, followed by an appropriate post-hoc test to compare individual groups. If the sample was not normally distributed, a Kruskal-Wallis test combined with Dunn’s post hoc test was used. For all statistical tests, differences were considered significant at p<0.05.

## Data and code availability

Scripts for the analysis of the clonality data are available through the lead author.

## Figure legends

**Figure S1: No reduction in OPC numbers following Nocodazole treatment**

**(A)** Experimental outline. GF = growth factors. **(B)** Number of OLIG2+ cells per well after treatment with nocodazole. Dashed line indicates the number of cells seeded at the beginning of the experiment. Related to Figure 2.

**Figure S2: Majority of OPCs are in G0/G1 after cell isolation**

**(A)** Percentage of living cells in different cell cycle phases. Related to Figure 2.

