## Supplementary figures and images for "Proliferation is a requirement for differentiation of oligodendrocyte progenitor cells during CNS remyelination"

### Suppl Fig 1

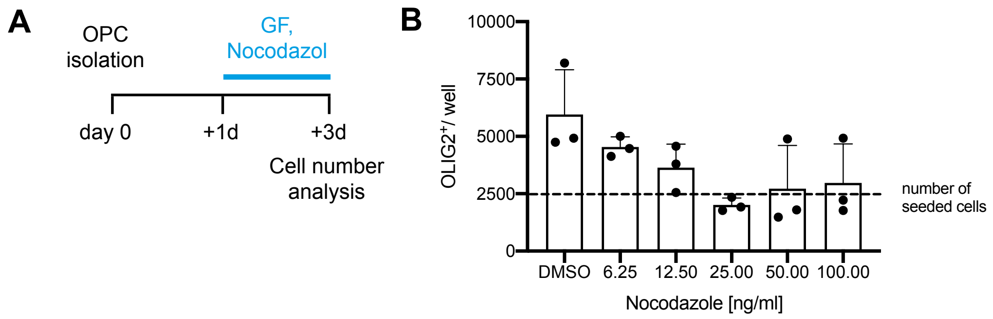


**Supplementary figure 1**

### Suppl Fig 2

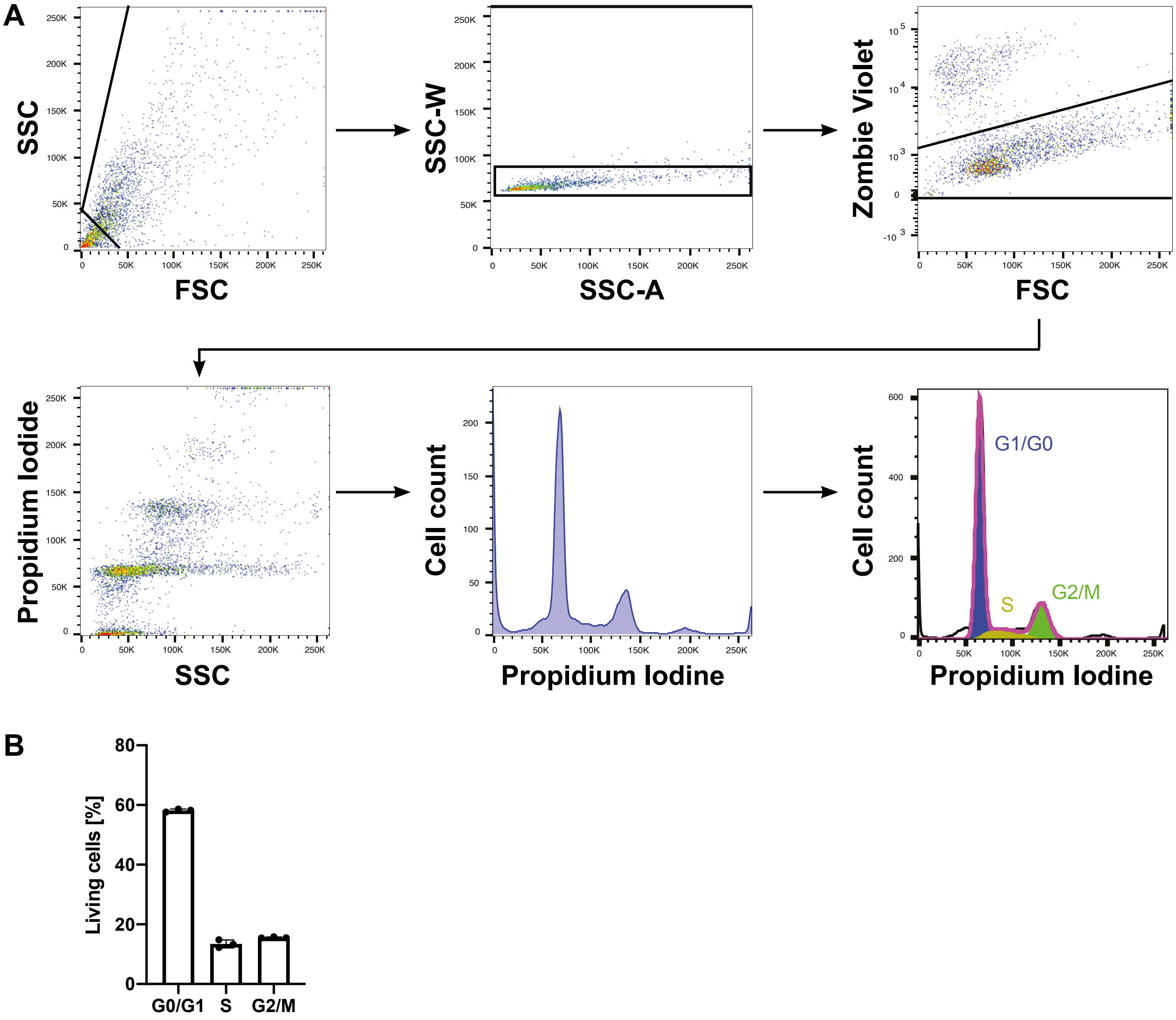


**Supplementary figure 2**
